# SARS-CoV-2 Neutralizing Antibody Responses Are More Robust in Patients with Severe Disease

**DOI:** 10.1101/2020.06.13.150250

**Authors:** Pengfei Wang, Lihong Liu, Manoj S. Nair, Michael T. Yin, Yang Luo, Qian Wang, Ting Yuan, Kanako Mori, Axel Guzman Solis, Masahiro Yamashita, Lawrence J. Purpura, Justin C. Laracy, Jian Yu, Joseph Sodroski, Yaoxing Huang, David D. Ho

## Abstract

We studied plasma antibody responses of 35 patients about 1 month after SARS-CoV-2 infection. Titers of antibodies binding to the viral nucleocapsid and spike proteins were significantly higher in patients with severe disease. Likewise, mean antibody neutralization titers against SARS-CoV-2 pseudovirus and live virus were higher in the sicker patients, by ~5-fold and ~7-fold, respectively. These findings have important implications for those pursuing plasma therapy, isolation of neutralizing monoclonal antibodies, and determinants of immunity.

---

The coronavirus disease 2019 (COVID-19), declared a pandemic by the World Health Organization, is an infection caused by a newly discovered coronavirus, severe acute respiratory syndrome coronavirus 2 (SARS-CoV-2)^1^. Together with SARS-CoV, which caused an outbreak 17 years ago, SARS-CoV-2 is a member of the subgenus *Sarbecovirus*. Both viruses express a glycoprotein termed spike protein (S), which mediates viral entry into ACE2-positive host cells and is therefore the target of virus-neutralizing antibodies^2–4^. Another structural protein is the nucleocapsid protein (NP), which is the most abundant and highly immunogenic protein in coronaviruses, making it a suitable candidate for diagnostic assays^5^.

A study of antibody responses to SARS-CoV-2 in patients with COVID-19 showed that nearly all patients developed virus-specific antibodies within 2-3 weeks after symptom onset^6^. Most serologic studies^6–9^ largely focused on binding antibodies to S and NP, but not virus-neutralizing antibodies even though such antibodies can be used therapeutically or prophylactically. Infusion of convalescent plasma has been used to treat SARS-CoV^10^ and SARS-CoV-2^11^. The measurement of neutralizing antibodies is critical in finding the best donors for plasma therapy, as well as being the gold standard to evaluate vaccine responses. Recent vaccine and re-infection studies in non-human primates suggest that neutralizing antibodies are the correlate of protection against SARS-CoV-2^12–14^. Studies using convalescent plasma to treat SARS-CoV-2 infections were performed only on a limited number of patients, and there were no careful measurements of neutralizing antibody titers to correlate with the clinical outcome^11,15–17^. This study examines SARS-CoV-2 neutralizing antibodies in the plasma of patients with different disease severity.

We studied 35 patients seen at Columbia University Irving Medical Center with PCR-confirmed SARS-CoV-2 infection to assess their plasma antibody responses to the virus. The age, sex, and time of blood collection after onset of symptoms for each patient are summarized in Extended Data Table 1. Patients who required hospitalization in the intensive care unit (19) were categorized as Severe, whereas those with milder disease with or without hospitalization (16) were categorized as Non-severe. As expected, Severe cases were older (range 34-84; mean 58) than Non-severe cases (range 20-58; mean 38). The sexes were equally distributed between both groups. Importantly, blood collection was taken, on average, about one month after the onset of symptoms in both groups (Extended Data Table 1).

Immunoassays to quantify antibodies to SARS-CoV-2 NP and S trimer, as well as to SARS-CoV S trimer were first developed (see Methods). These assays were next evaluated using five normal serum samples plus serum samples from two patients who recovered from PCR-confirmed SARS-CoV-2 infection. Indeed, the normal sera lacked detectable binding antibodies to the viral proteins, whereas the two convalescent sera showed robust binding antibody titers to SARS-CoV-2 NP and S trimer as well as modest cross-reactivity to SARS-CoV S trimer (Extended Data Fig. 1). These antibody binding assays were then used to measure binding antibody titers in plasma of both Severe and Non-severe patients. Plasma titers of antibodies to SARS-CoV-2 NP and S trimer were substantially higher in Severe patients than in Non-severe patients (Fig. 1a, left and middle panels). Specifically, for NP-directed antibodies, the reciprocal plasma titers ranged from 292 to 37,099 (mean 5,086) for Severe cases and from 170 to 1,376 (mean 615) for Non-severe cases (Fig. 1b). The mean plasma titer was ~8-fold higher in the Severe group, and this difference was statistically significant (p = 0.036). Similarly, for S trimer-directed antibodies, the reciprocal plasma titers ranged from 257 to 18,397 (mean 2,985) for Severe patients and from <100 to 1,963 (mean 364) for Non-severe patients (Fig. 1b and Extended Data Table 1). The mean plasma titer was also ~8-fold higher in the Severe group, and this difference was again statistically significant (p = 0.016). A strong positive correlation was observed between NP antibody titers and S trimer antibody titers (Extended Data Fig. 2). Cross-reactive antibody titers to SARS-CoV S trimer were lower overall, but still more robust in patients with severe disease (Fig. 1, right panels). Only one plasma sample, S2, had a high titer to SARS-CoV S trimer.

**Fig. 1.**
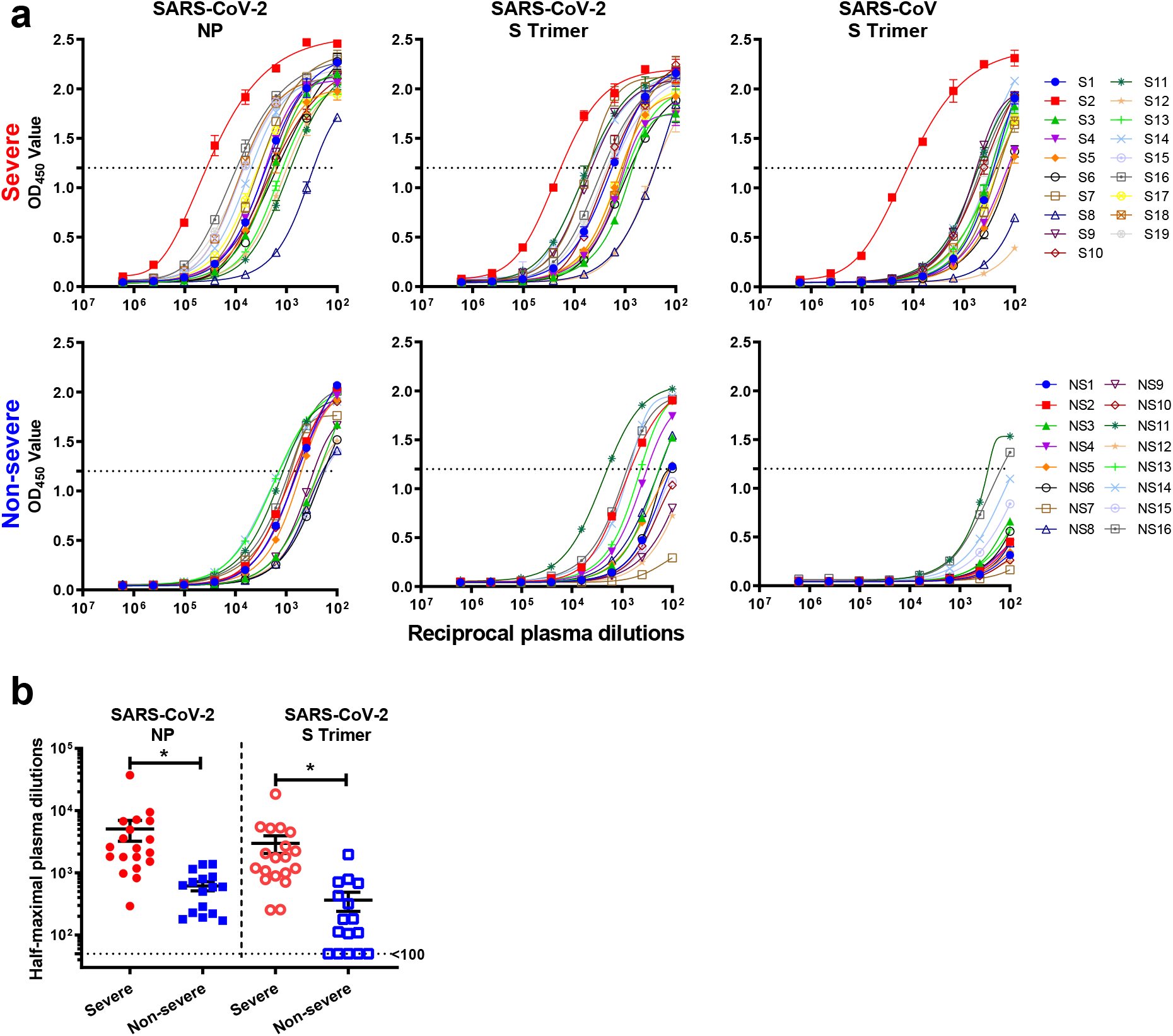
Binding antibody responses against SARS-CoV-2 and SARS-CoV. **a**, Plasma titers of binding antibodies to SARS-CoV-2 NP, S trimer, and SARS-CoV S trimer in Severe and Non-severe patients. **b**, Comparison of the level of binding antibodies against SARS-CoV-2 between Severe and Non-severe patients. Lines represent mean ± SEM and *p* values were calculated by two-tailed *t*-test. *, *p* < 0.05.

Next, antibody neutralization assays against SARS-CoV-2 pseudovirus and live virus (2019-nCoV/USA_WA1/2020) were developed (see Methods). These assays were then evaluated using the aforementioned five normal serum samples and two serum samples from PCR-confirmed cases of SARS-CoV-2 infection. Convalescent sera exhibited robust neutralizing antibody titers against both the pseudovirus and the live virus, whereas the five normal sera exhibited none (Extended Data Fig. 3). There was no significant cross-neutralization of the SARS-CoV pseudovirus by the convalescent sera. All plasma samples from Severe and Non-severe patients were then tested to determine their virus-neutralizing titers. Overall, both SARS-CoV-2 pseudovirus and live virus neutralization titers were substantially higher in the plasma of Severe patients compared to those of Non-severe patients (Fig. 2, left and middle panels). Specifically, in the SARS-CoV-2 pseudovirus assay, the reciprocal plasma neutralizing titers ranged from <100 to 13,710 (mean 2,545) for Severe cases and from <100 to 1,463 (mean 491) for Non-severe cases (Fig. 2b and Extended Data Table 1). The mean pseudovirus neutralizing titer was ~5-fold higher in the Severe group, and this difference was statistically significant (p = 0.015). Similarly, in the SARS-CoV-2 live virus assay, the reciprocal plasma titers ranged from 926 to 30,175 (mean 10,701) for Severe patients and from <100 to 6,884 (mean 1,485) for Non-severe patients (Fig. 2b and Extended Data Table 1). The mean live virus neutralizing titer was ~7-fold higher in the Severe group, and this difference was again statistically significant (p < 0.001). Only plasma samples from Severe patients show cross-neutralization against SARS-CoV pseudovirus (Fig. 2, right panels), albeit at much lower titers.

**Fig. 2.**
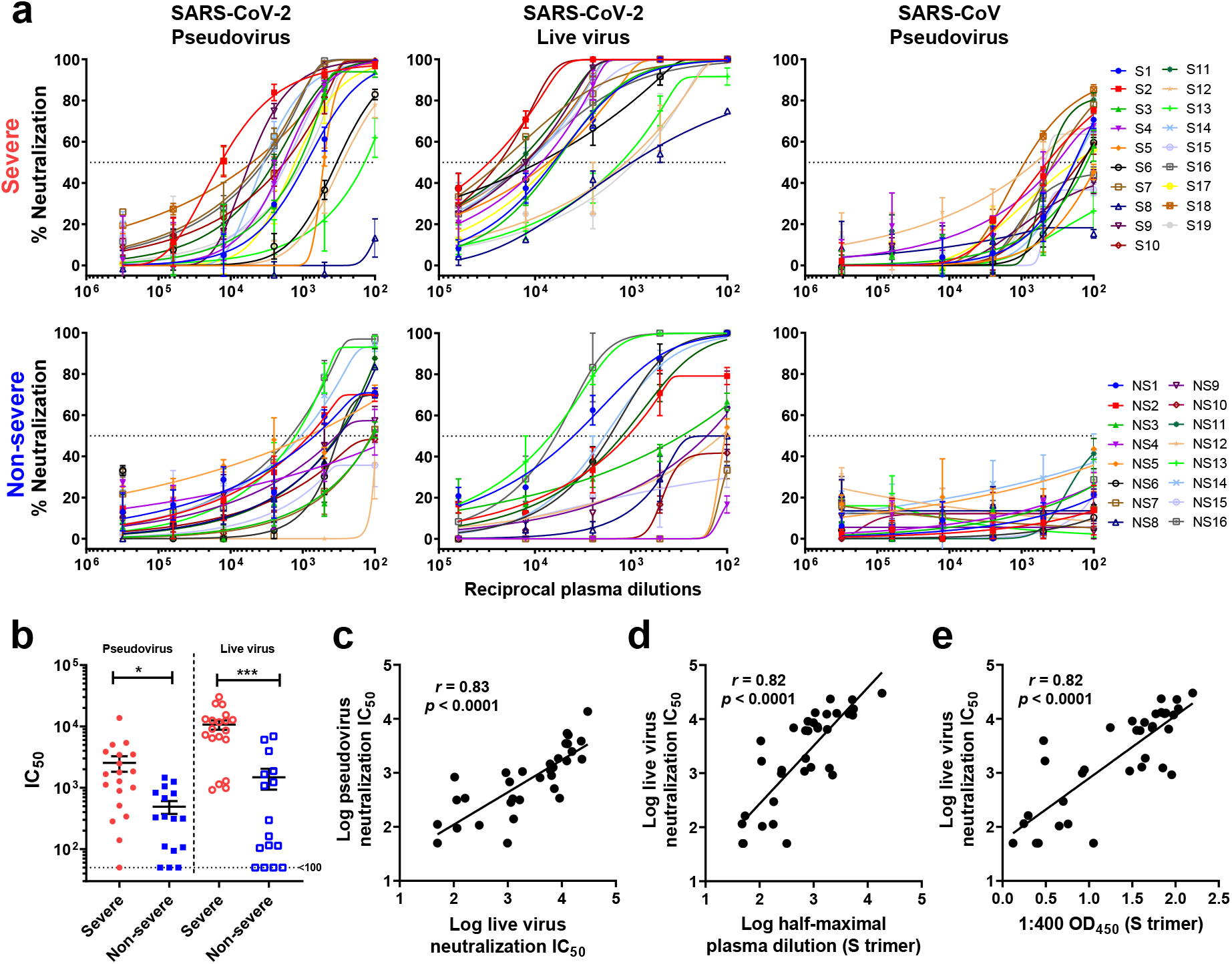
Neutralizing antibody responses against SARS-CoV-2 and SARS-CoV. **a**, Plasma neutralizing activities against SARS-CoV-2 pseudovirus, live virus and SARS-CoV pseudovirus in Severe and Non-severe patients. **b**, Comparison of the level of neutralizing antibodies against SARS-CoV-2 between Severe and Non-severe patients. **c-e**, Correlation of SARS-CoV-2 live virus neutralization tiers versus pseudovirus neutralization titers (**c**), S trimer-binding antibody titers (**d**) and OD450 values at 1:400 plasma dilution (**e**). Lines in **b** represent mean ± SEM and *p* values were calculated by two-tailed *t*-test. *, *p* < 0.05; ***, *p* < 0.001. In **c-e**, the *Pearson correlation coefficient (r)* and the probability *p* value were calculated using GraphPad Prism.

A few other findings were notable. First, the plasma neutralizing titers against the SARS-CoV-2 pseudovirus correlated quite well with the titers obtained against the live virus (Fig. 2c). In addition, neutralizing titers against the pseudovirus (Extended Data Fig. 4) or live virus (Fig. 2d) correlated well with S trimer-binding antibody titers as determined by quantitative immunoassay. In fact, even an optical density reading (OD_450_) at one plasma dilution of 1:400 correlated well with virus neutralization titers (Extended Data Fig. 4 and Fig. 2e).

The results of this study show that patients with severe SARS-CoV-2 disease have more robust binding antibodies to both NP and S trimer (Figs. 1a and 1b). Functionally active antibodies capable of virus neutralization were also more abundant in the sicker patients (Figs. 2a and 2b). The latter finding is reminiscent of the observation that HIV-1 broadly neutralizing antibodies were most commonly found in patients with persistent viremia for a protracted period^18^. There is evidence that patients with severe SARS-CoV-2 infection have a higher viral load^9^, and perhaps a longer exposure to a greater abundance of viral antigens is the basis for our findings. Regardless, the results reported herein do have important implications for donor selection when pursuing plasma therapy^19^ or isolating neutralizing monoclonal antibodies^20^. Of course, this selection is best made by assessing virus-neutralizing activity in the serum or plasma of potential donors. However, even an S trimer-based immunoassay could provide useful guidance in choosing convalescent patients who have the most robust neutralizing antibodies (Figs. 2d and 2e; Extended Data Fig. 4). The scientific community and general public eagerly await data that could answer whether having virus-neutralizing antibodies is equivalent to having protective immunity. The strong correlations observed here between antibodies that bind the S trimer and antibodies that neutralize the pseudovirus (Extended Data Fig. 4) or live virus (Figs. 2d and 2e) could facilitate future studies to understand what constitutes immunity against SARS-CoV-2.

