## Supplementary material for "SARS-CoV-2 Neutralizing Antibody Responses Are More Robust in Patients with Severe Disease": Methods and Extended Data

**Study design, population, and serum samples.** This was a cross-sectional study of SARS-CoV-2 neutralizing antibody responses in participants enrolled in an observational cohort study of convalescent patients in the New York City area. Eligibility criteria included a diagnosis of SARS-CoV-2 infection by RT-PCR prior to enrollment, without restriction on duration of time since symptom onset. Plasma obtained at the enrollment visit from participants with severe (requiring intensive care unit monitoring and mechanical ventilation) and non-severe clinical presentations were analyzed for SARS-CoV-2 antibody responses.

**Protection of human subjects and confidentiality.** This observational cohort study was approved by the Institutional Review Board (IRB) of the Columbia University Irving Medical Center and all participants provided written informed consent.

**Recombinant protein expression.** Spike trimer and nucleocapsid were expressed and purified as previously described[21]. Briefly, sequences corresponding to the ectodomains of the spike (S) for SARS-CoV-2 and SARS-CoV were cloned into mammalian expression vectors pCAGGS. For each construct, a foldon trimerization domain followed by a 6x His tag and a StrepTag II® were fused to the C-terminus of the gene. These constructs were transiently transfected into Expi293 cells and then the expressed protein was purified by using the Strep-Tactin XT Resin (cat# P2004-1-5, Zymo Research) as recommended by the manufacturer. The nucleocapsid protein (NP) of SARS-CoV-2 was cloned into a bacterial expression vector pET28a(+). A 6x His tag was placed at the C-terminus of NP. *E.coli* were transformed with the construct and then the expressed protein was purified by using Ni-NTA agarose (cat# K95001, Invitrogen) as recommended by the manufacturer, then followed by the purification on a HiTrap Heparin-HP column and on a Superdex 200 10/300 GL column[22].

**ELISA.** Wells of 96-well EIA/RIA Assay Microplates (Corning) were coated with 50 ng of SARS-CoV or SARS-CoV-2 spike trimer or SARS-CoV-2 NP overnight at 4 °C. After washing with 0.05% Tween-20 in PBS (PBST), plates were blocked with 300 μL/well of blocking buffer (1% BSA and 10% bovine calf serum in PBS) for 1 h at 37 °C, then washed again with PBST. Antibodies or heat-inactivated plasma samples from COVID-19 patients or healthy donors were serially diluted in dilution buffer (1% BSA and 20% bovine calf serum in PBS) and then incubated in the plates for 1 h at 37 °C. Afterwards, plates were washed with PBST and then incubated with Peroxidase AffiniPure goat anti-human IgG (H+L) and goat anti-human IgM (cat# 109-035-003 and 109-035-043, Jackson ImmunoResearch, both at 1:10,000 dilution) antibodies for 1 h at 37 °C. Plates underwent a final wash with PBST and then the antibody binding was detected by incubating with TMB substrate (cat# 4444, Sigma) for 3 min. The reaction was stopped by adding 1N sulfuric acid (cat# SA212-1, Fisher Scientific). Absorbance was measured at 450 nm and the OD450 values were analyzed by using GraphPad Prism 8 (GraphPad Software, Inc.).

**Production of pseudoviruses.** Recombinant Indiana VSV (rVSV) expressing SARS-CoV-2 and SARS-CoV spikes were generated as previously described[23,24]. HEK293T cells were grown to 80% confluency before transfection with pCMV3-SARS-CoV-2-spike or pCMV3-SARS-CoV-spike (kindly provided by Dr. Peihui Wang, Shandong University, China) using FuGENE 6 (Promega). Cells were cultured overnight at 37°C with 5% $CO_2$. The next day, medium was removed and VSV-G pseudotyped ΔG-luciferase (G*ΔG-luciferase, Kerafast) was used to infect the cells in DMEM at an MOI of 3 for 1 hr before washing the cells with 1X DPBS three times. DMEM supplemented with 2% fetal bovine serum and 100 I.U./mL penicillin and 100 μg/mL streptomycin was added to the infected cells and they were cultured overnight as described above. The next day, the supernatant was harvested and clarified by centrifugation at 300g for 10 min before aliquoting and storing at −80°C.

**Pseudovirus-based neutralization assay**. Neutralization assays were performed by incubating pseudoviruses with serial dilutions of heat-inactivated plasma and measured by the reduction in luciferase gene expression. In brief, Vero E6 cells (ATCC) were seeded in a 96-well plate at a concentration of $2 \times 10^4$ cells per well. The next day, pseudoviruses were incubated with serial dilutions of the plasma samples (six dilutions in a 5-fold step-wise manner) in duplicate for 30 min at 37°C. The mixture was added to cultured cells for infection and incubated for an additional 24 hrs. The luminescence was measured by Britelite plus Reporter Gene Assay System (PerkinElmer). The 50% inhibitory dilution ($IC_{50}$) was defined as the plasma dilution at which the relative light units (RLUs) were reduced by 50% compared with the virus control wells (virus + cells) after subtraction of the background RLUs in the control groups with cells only. The $IC_{50}$ values were calculated with non-linear regression using GraphPad Prism 8.

**Neutralization of SARS-CoV-2 live virus.** We used an end-point dilution in a 96-well plate format to measure the neutralization of SARS-CoV-2. The infectious isolate (2019-nCoV/USA_WA1/2020) for these experiments was provided by the World Reference Center for Emerging Viruses and Arboviruses located at University of Texas Medical Branch, Galveston, TX. In brief, patient plasma samples were heat-inactivated by incubation at 56°C for 30 minutes and serially diluted at 5-fold dilutions starting at a 1:100 dilution. Each dilution of the plasma was tested in triplicates. Diluted samples were incubated with SARS-CoV-2 at an MOI of 0.2 in EMEM with 7.5% inactivated fetal calf serum for 1 hour at 37°C. Post incubation, the virus-plasma mix was transferred on a 24hr-old monolayer of Vero E6 cells containing about 60,000 cells in 100µL of E10 culture medium. The cells were incubated with the mixture for 70 hours. Morphological changes resulting from cytopathic effects (CPE) of the virus were visually scored for each well from 1-4, with 4 being the appearance of complete virus-mediated cytopathy. Double-blinded scoring of the cytopathic effect was converted to percentage of neutralization and the $IC_{50}$ calculated with non-linear regression using GraphPad Prism 8.

**Data availability**

All of the data supporting this study are shown in the main text figures and Extended Data. The materials involved in this study are available upon request to the corresponding author (D.D.H).

**Acknowledgements**

We thank Professor Kwok-yung YUEN, University of Hong Kong and Eldad A. Hod, Columbia University for providing some of the patient samples. This work was supported by the Jack Ma Foundation, The JBP Foundation, and the Wu Family China Center.

**Author contributions**

Conceptualization was provided by D.D.H. The methodology was developed by P.W., L.L., M.N., and Y.H. Investigations were carried out by P.W., L.L., M.N., Y.H., Y.L., Q.W., T.Y., K.M., A.G.S., M.Y., J.Y., J.S., Resources were provided by M.T.Y, L.J.P, and J.C.L. The manuscript was written by D.D.H., P.W., L.L., M.N., and M.T.Y., and reviewed and commented on by all the authors. Y.H. and D.D.H. provided supervision.

**Competing interests**

The authors declare no competing interests.

### Additional information

Extended data is available for this paper at

### Extended Data Table 1

| Sample ID | Age | Sex | DASO[a] | S Trimer ELISA | | Neutralization $IC_{50}$ | |
|---|---|---|---|---|---|---|---|
| | | | | 1: 400 $OD_{450}$ | Half-maximal plasma dilution | Pseudovirus | Live virus |
| S1[b] | 40 | M | 24 | 1.92 | 1,791 | 893 | 6,441 |
| S2 | 46 | M | 30 | 2.20 | 18,397 | 13,710 | 30,175 |
| S3 | 65 | F | 28 | 1.55 | 708 | 1,121 | 6,140 |
| S4 | 71 | F | 45 | 1.64 | 993 | 1,846 | 8,578 |
| S5 | 61 | M | 37 | 1.74 | 1,175 | 511 | 7,263 |
| S6 | 62 | F | 36 | 1.50 | 782 | 339 | 9,113 |
| S7 | 38 | M | 25 | 2.02 | 5,198 | 3,864 | 22,898 |
| S8 | 62 | F | 33 | 0.92 | 253 | <100 | 987 |
| S9 | 59 | M | 37 | 1.87 | 4,518 | 5,404 | 12,500 |
| S10 | 34 | M | 28 | 1.84 | 2,039 | 1,784 | 23,585 |
| S11 | 46 | M | 30 | 2.03 | 5,476 | 2,485 | 15,392 |
| S12 | 45 | F | 32 | 0.95 | 257 | 284 | 1,140 |
| S13 | 53 | M | 24 | 1.61 | 893 | 140 | 1,282 |
| S14 | 66 | F | 26 | 1.98 | 5,293 | 3,497 | 11,797 |
| S15 | 78 | F | 30 | 1.77 | 1,748 | 1,666 | 12,358 |
| S16 | 57 | F | 37 | 1.89 | 2,683 | 3,447 | 12,796 |
| S17 | 76 | M | 31 | 1.73 | 1,195 | 1,279 | 6,807 |
| S18 | 50 | M | 19 | 1.84 | 1,073 | 5,033 | 13,136 |
| S19 | 84 | F | 32 | 1.96 | 2,238 | 1,006 | 926 |
| **Mean** | **58** | | **31** | **1.73** | **2985** | **2545** | **10701** |
| **Median** | **59** | | **30** | **1.84** | **1748** | **1666** | **9113** |
| NS1[c] | 44 | M | 27 | 0.47 | 106 | 812 | 3,960 |
| NS2 | 38 | F | 28 | 1.47 | 679 | 681 | 1,081 |
| NS3 | 20 | M | 36 | 0.70 | 182 | 107 | 297 |
| NS4 | 50 | F | 30 | 1.05 | 318 | <100 | <100 |
| NS5 | 50 | M | 30 | 0.66 | 112 | 834 | 104 |
| NS6 | 37 | F | 32 | 0.49 | 109 | 315 | 1,662 |
| NS7 | 25 | M | 35 | 0.12 | <100 | 111 | <100 |
| NS8 | 40 | M | 25 | 0.76 | 179 | 313 | 113 |
| NS9 | 32 | F | 31 | 0.30 | <100 | 338 | 162 |
| NS10 | 58 | F | 34 | 0.42 | <100 | <100 | <100 |
| NS11 | 49 | M | 35 | 1.85 | 1,963 | 329 | 1,235 |
| NS12 | 25 | F | 27 | 0.25 | <100 | <100 | 115 |
| NS13 | 32 | M | 34 | 1.25 | 427 | 1,254 | 6,884 |
| NS14 | 34 | M | 35 | 1.65 | 711 | 1,057 | 1,875 |
| NS15 | 31 | F | 35 | 0.39 | <100 | <100 | <100 |
| NS16 | 50 | F | 36 | 1.59 | 789 | 1,463 | 6,063 |
| **Mean** | **38** | | **32** | **0.84** | **364** | **491** | **1485** |
| **Median** | **38** | | **33** | **0.68** | **146** | **322** | **230** |
| ***p* value** | **0.0001** | | **0.5151** | **<0.0001** | **0.0164** | **0.0147** | **0.0001** |

[a]Days after symptom onset, [b]Severe, [c]Non-severe

**Table Footnote:** In the **Severe Group (N=19)**, mean age was 58, 58% were male. All received mechanical ventilation in the intensive care unit, 6 (32%) had acute kidney injury requiring dialysis, and received the following treatment for COVID-19 during their hospital course: hydroxychloroquine (89%), remdesivir (16%), and tocilizumab (11%). At the time of manuscript writing, 4 (21%) died during their hospitalization, 5 (26%) remain hospitalized with a mean duration of 75 days, and 11 (58%) have been discharged from the hospital.

In the **Non-Severe Group (N=16),** mean age was 38, 56% were male, and only one (NS11) required hospitalization, but not intensive care unit monitoring or mechanical ventilation.

The value for the **Severe Group** and **Non-Severe Group** were compared by two-tailed *t*-test; the *p* values for these comparisons are shown.

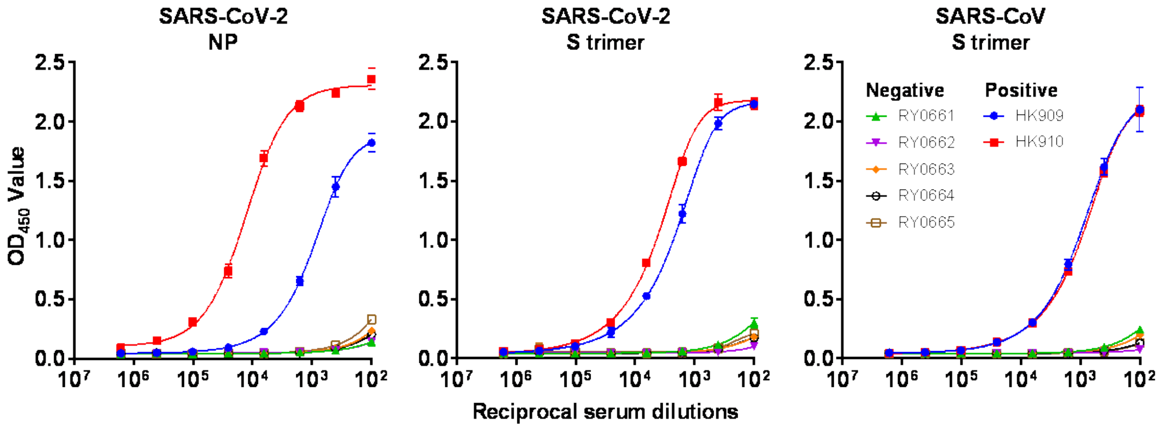

**Extended Data Fig. 1 | ELISA validation.** ELISAs detecting binding antibodies to SARS-CoV-2 NP, S trimer, and SARS-CoV S trimer were evaluated using five normal serum samples (RY0661-RY0665) plus serum samples from two patients who recovered from PCR-confirmed SARS-CoV-2 infection (HK909 and HK910).

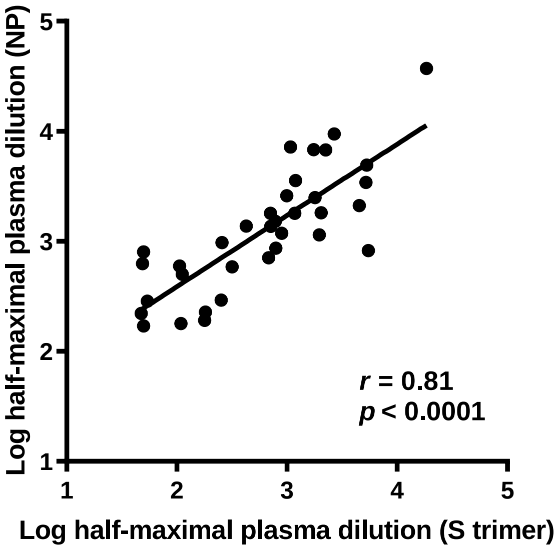

**Extended Data Fig. 2 | Correlation of SARS-CoV-2 S trimer-binding antibody titers versus NP-binding antibody titers.** The *Pearson correlation coefficient (r)* and the probability *p* value were calculated using GraphPad Prism.

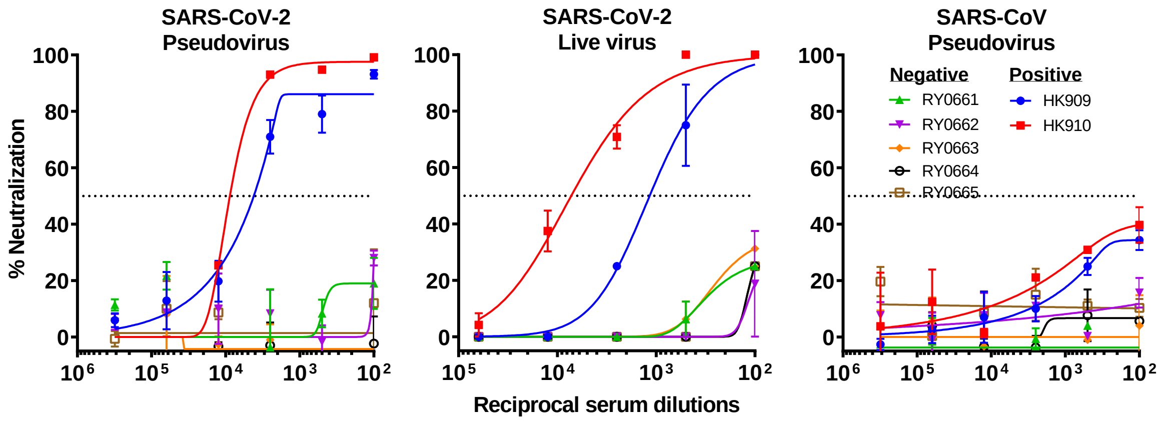

**Extended Data Fig. 3 | Neutralization assays validation.** The assays detecting neutralizing antibodies agianst SARS-CoV-2 pseudovirus, live virus and SARS-CoV pseudovirus were evaluated using five normal serum samples (RY0661-RY0665) plus serum samples from two patients who recovered from PCR-confirmed SARS-CoV-2 infection (HK909 and HK910).

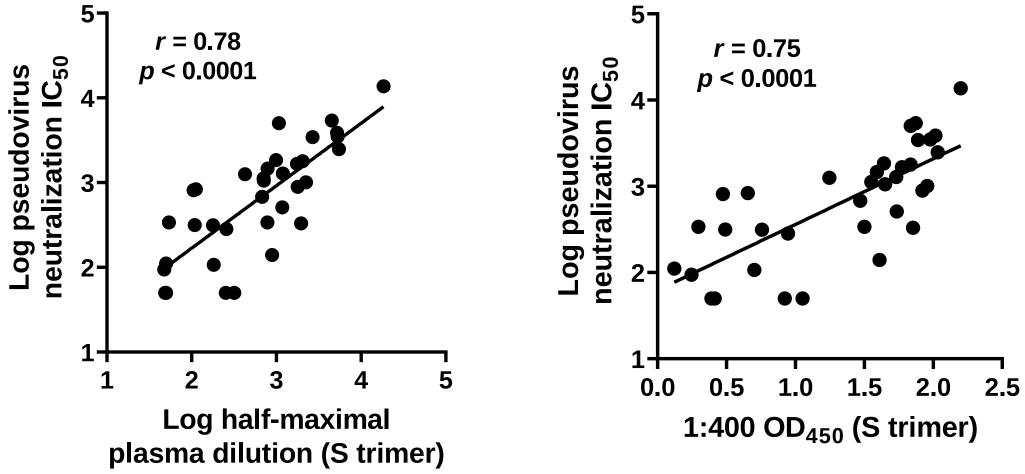

**Extended Data Fig. 4 | Correlation of SARS-CoV-2 pseudovirus neutralization titers versus S trimer-binding antibody titers (left) and $OD_{450}$ values at 1:400 plasma dilution (right).** The *Pearson correlation coefficient (r)* and the probability *p* value were calculated using GraphPad Prism.
